# Multimodal Assessment of Non-Alcoholic Fatty Liver Disease with Transmission-Reflection Optoacoustic Ultrasound

**DOI:** 10.1101/2022.08.16.504139

**Authors:** Berkan Lafci, Anna Hadjihambi, Christos Konstantinou, Joaquin L. Herraiz, Luc Pellerin, Neal C. Burton, Xosé Luís Deán-Ben, Daniel Razansky

## Abstract

Non-alcoholic fatty liver disease (NAFLD) is an umbrella term referring to a group of conditions associated to fat deposition and damage of liver tissue. Early detection of fat accumulation is essential to avoid progression of NAFLD to serious pathological stages such as liver cirrhosis and hepatocellular carcinoma. We exploited the unique capabilities of transmission-reflection optoacoustic ultrasound (TROPUS), which combines the advantages of optical and acoustic contrasts, for an early-stage multi-parametric assessment of NAFLD in mice. The multispectral optoacoustic imaging allowed for spectroscopic differentiation of lipid content, as well as the bio-distributions of oxygenated and deoxygenated hemoglobin in liver tissues *in vivo*. The pulse-echo (reflection) ultrasound (US) imaging further provided a valuable anatomical reference whilst transmission US facilitated the mapping of speed of sound changes in lipid-rich regions, which was consistent with the presence of macrovesicular hepatic steatosis in the NAFLD livers examined with *ex vivo* histological staining. The proposed multimodal approach facilitates quantification of liver abnormalities at early stages using a variety of optical and acoustic contrasts, laying the ground for translating the TROPUS approach toward diagnosis and monitoring NAFLD in patients.

## I. INTRODUCTION

Non-alcoholic fatty liver disease (NAFLD) is a common disorder comprising a progressive spectrum of diseases, defined as an accumulation of fat in the liver (steatosis), in the absence of significant alcohol consumption [1]. NAFLD progresses to non-alcoholic steatohepatitis, characterized by inflammation and hepatocyte damage (which includes ballooning and cell death), together with deposition of collagen and fibrosis progression [2], results in enlarging and discoloring of the organ [3]. Further progression of fibrosis may lead to the irreversible stages of cirrhosis and, eventually, hepatocellular carcinoma [4]. At present, no approved interventions are available to treat liver fibrosis, which calls for the development of new research tools aimed at better understanding the underlying causes of NAFLD, as well as new methods capable of detecting this condition at the earliest reversible stage before it progresses to fibrosis [5], [6]. NAFLD and liver fibrosis have become a major health concern due to the growing prevalence of overweight and obese individuals in developed countries [7]. The worldwide mortality rate related to liver diseases follows an upward trend, reaching 2 million disease-related deaths annually in 2019 [8]. However, detection of early liver damage is challenged by the small size and sparsity of the scars formed before the appearance of fibrosis [9]. Currently, liver disease assessment is performed with biopsies and histopathology imaging [10]. Liver biopsy is however an invasive and user-dependent (sampling bias) procedure hindering a continuous monitoring of liver tissue abnormalities. Therefore, the development of non-invasive methods enabling the quantitative assessment of NAFLD is paramount both for preclinical studies aiming at advancing our knowledge of the disease, as well as for early diagnosis purposes in the clinical setting.

Whole-body clinical imaging methods have been shown to provide important advantages for liver disease diagnosis. Magnetic resonance imaging (MRI) achieves high specificity for fat accumulation by using the proton density fat fraction technique [11]. X-ray computed tomography (CT) has also been reported for the assessment of liver abnormalities with high resolution [12]. However, the use of these methods is associated with high installation and maintenance costs, exposure to ionizing radiation, and insufficient sensitivity to molecular (fat) contrast [13], [14]. Ultrasound (US) imaging is a more affordable and accessible bedside technology which has also been used for visualizing and assessing liver abnormalities [15], [16]. Linear array probes are typically used in clinics to provide a quick assessment of the liver with pulse-echo (B-mode) US. However, this approach does not provide sufficient tomographic (angular) coverage needed for accurate localization and quantitative characterization of the damaged liver areas, further lacking the necessary contrast for assessing fat accumulation. In response, tomographic US methods have been developed to provide enhanced tissue contrast. Reflection ultrasound computed tomography (RUCT) is based on tomographic pulse-echo US imaging with waves being sequentially emitted and detected at different angular positions around the sample. The broad angular coverage has been shown to increase the image contrast, resolution and field of view (FOV) with respect to those achieved with linear arrays [17]. Transmission ultrasound computed tomography (TUCT) further enables mapping the speed of sound (SoS) distribution in tissues by considering US waves transmitted through the sample. SoS maps have been shown to provide improved specificity for detecting fatty and glandular tissue abnormalities and delineation of lesions [18].

Hybrid optoacoustic (OA) imaging combining light with sound has emerged as another powerful functional and molecular preclinical imaging approach. It is based on optical excitation of tissues at near-infrared (NIR) wavelengths and tomographic detection of the thermoelastically-induced US waves, thus rendering rich optical contrast with high spatial resolution unaffected by photon scattering in deep tissues [19], [20]. In particular, multispectral optoacoustic tomography (MSOT) capitalizes on optical excitation at different wavelengths to spectroscopically differentiate between oxy- (HbO_2_) and deoxy-hemoglobin (Hb), melanin, lipids and other tissue bio-chromes as well as extrinsically administered contrast agents [21],[22]. However, unambiguous anatomical differentiation of lesions and organs is hindered with MSOT whose main contrast stems from hemoglobin-rich structures such as major blood vessels.

Recently, a multi-modal transmission-reflection optoacoustic ultrasound (TROPUS) imaging has been suggested as a versatile imaging approach for multi-parametric anatomical, functional and molecular characterization of murine disease models [23], [24]. The full tomographic coverage of the circular transducer array used in TROPUS results in an improved contrast and resolution with MSOT, RUCT and TUCT, while further providing real-time imaging capabilities for visualizing dynamic processes [25], [26]. Here, we employed TROPUS for assessing early-stage NAFLD in mice. The lipid accumulation in the liver was delineated and quantified with TUCT and MSOT while RUCT facilitated anatomical interpretation. The *in vivo* imaging results were validated with Haematoxylin and Eosin (H&E) staining of excised specimens.

## II. RESULTS

The TROPUS imaging setup consists of a circular US transducer array, a nanosecond laser source, a data acquisition-transmission system (DAQ) and a workstation PC used for the system synchronization, data transfer, storage and processing (Fig. 1a, see Methods for details) [24]. Imaging in the MSOT mode was performed by quickly switching the optical wavelength of the nanosecond optical parametric oscillator (OPO) laser from 740 to 940 nm with a 20 nm step size at 25 Hz repetition rate in order to enable the separation of Hb, HbO_2_, melanin and lipid components, the latter having a distinct peak in its absorption spectrum at 920 nm (Fig. 1b) [27]. Exemplary cross-sectional MSOT images acquired from living mice at 800nm excitation wavelength are shown in Fig. 1c. RUCT imaging was based on the synthetic transmit aperture (STA) image acquisition technique [25], which employs sequential transmission of US pulses with each array element followed by detection of the reflected signals. Image compounding was subsequently performed by adding up multiple low-contrast delay-and-sum images acquired from different views around the sample, resulting in a final high-contrast RUCT images (Fig. 1d). Quantitative TUCT images representing the SoS distribution in the mouse in m/s were reconstructed from the US waves that traversed the imaged object using a full wave inversion (FWI) algorithm (Fig. 1e) [28].

**Figure 1:**
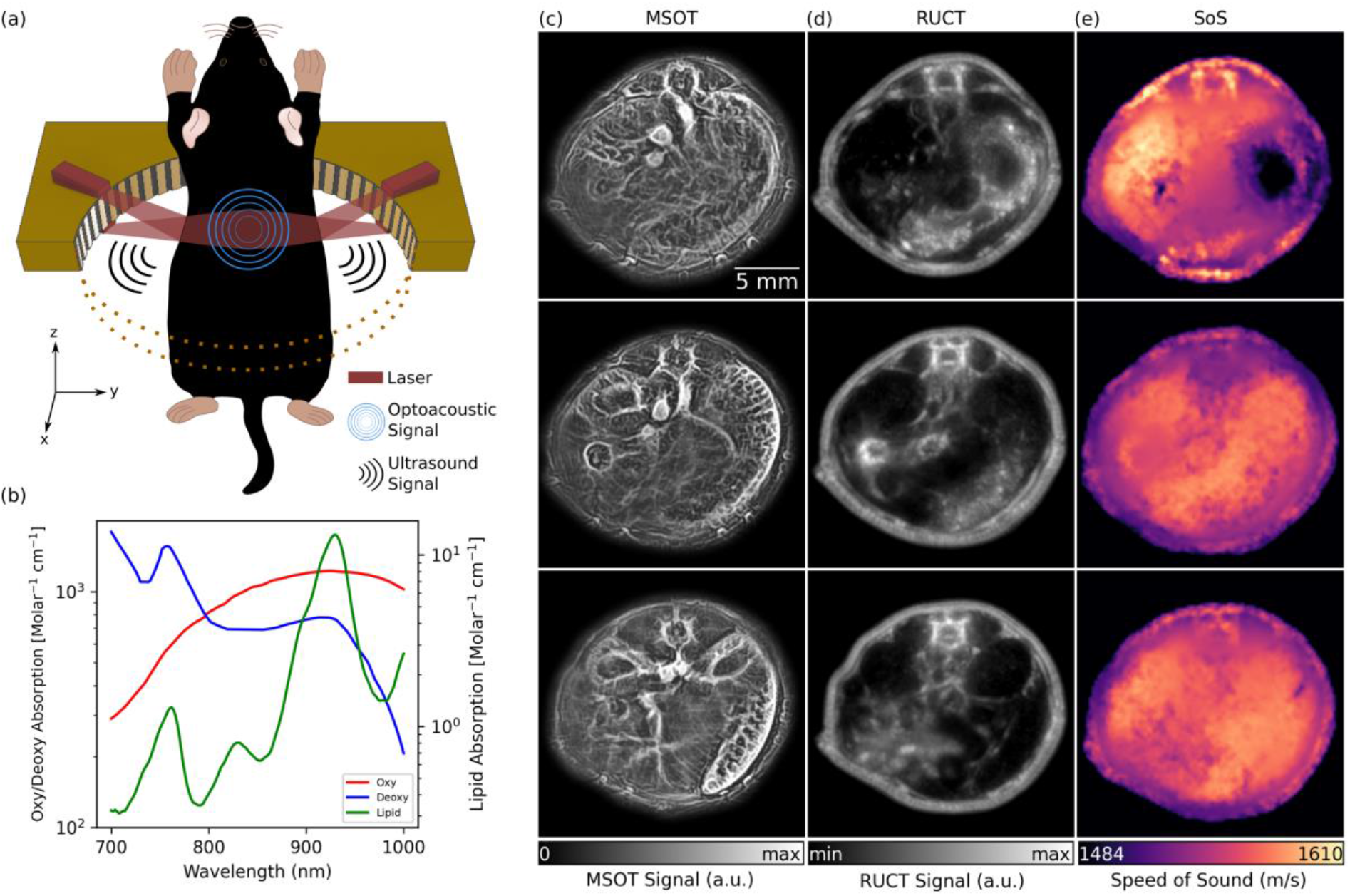
TROPUS imaging. (a) Lay-out of the imaging set-up combining three modalities, namely multispectral optoacoustic tomography (MSOT), reflection ultrasound computed tomography (RUCT) and speed of sound (SoS) imaging. Only half of the ring array is shown for better visualization. (b) Absorption spectrum of oxy-hemoglobin (HbO_2_), deoxy-hemoglobin (Hb) and lipid in 700 nm and 1000 nm wavelength range. (c) Exemplary MSOT images from different cross sections recorded at 1064 nm excitation wavelength. (d) The corresponding cross sections reconstructed with the RUCT modality. (e) The corresponding cross sections showing the SoS maps reconstructed with the transmission ultrasound computed tomography (TUCT) modality.

The basic ability of the multimodal TROPUS system to differentiate between NAFLD and control liver tissues was first evaluated with *ex vivo* samples (Fig. 2). Specifically, livers excised from 3 NAFLD and 3 control mice were imaged at two different vertical positions, resulting in 12 cross-sectional images. The MSOT images enable resolving the fat content by capitalizing on the distinctive optical absorption spectrum of lipids (Fig. 1b). The spectrally un-mixed bio-distributions of lipids (green color in Fig. 2a), overlaid onto the structural MSOT images rendered by averaging signals acquired at all the excitation wavelengths, clearly evince a higher fat content in the NAFLD liver tissue with respect to the control. RUCT images were further acquired for anatomical reference (Fig. 2b), which were used to delineate the borders of the excised livers in order to create binary masks to suppress background and conduct quantitative analysis. The corresponding SoS images acquired with TUCT manifest lower SoS values in the entire cross-sections of NAFLD livers as compared to the controls (Fig. 2c), which is generally expected considering a slower sound wave propagation in fat compared to liver tissue [29]. Histology images based on H&E staining were also acquired for validation (Fig. 2d). The spectra of the MSOT signals averaged over selected regions of interest (ROIs) revealed the presence of fat in the liver tissue from animals with NAFLD (Fig. 2e). Specifically, a distinctive peak at 920 nm was observed in the MSOT signal spectra matching well the known local maximum in the optical absorption of lipids [27]. This spectral peak was not present in the spectrum of the MSOT signals recorded from the control liver tissue. The lipid signals in *ex vivo* liver tissues were then averaged based on the pixel number after removing the non-distinct absorption background. The averaged lipid signal un-mixed from the MSOT images was 47% higher in NAFLD livers as compared to controls, exhibiting statistically significant differences for the 12 imaged cross-sections (Fig. 2f, p=0.001). Note that a similar standard deviation (STD) of the lipid signal (12% of the average value of all images) was observed in both cases. Statistical analysis of the measured SoS values for the 12 imaged cross-sections further revealed significant differences between NAFLD liver tissues and controls (Fig. 1g, p=0.010). The measured SoS mean and STD in NAFLD mice were 1495 m/s and 12 m/s, respectively, while these values were 1525 m/s and 15 m/s, respectively, in control mice. Histology images based on H&E staining further revealed macrovesicular hepatic steatosis in the NAFLD livers (Figs. 2d). The lipid accumulates in the hepatocytes as vacuoles detectable with H&E staining [30]. These intracytoplasmatic fat droplets were not observed in histology images of control tissues. The difference in fat content observed in histology images is thus consistent with the observations in the *in vivo* MSOT and SoS images.

**Figure 2:**
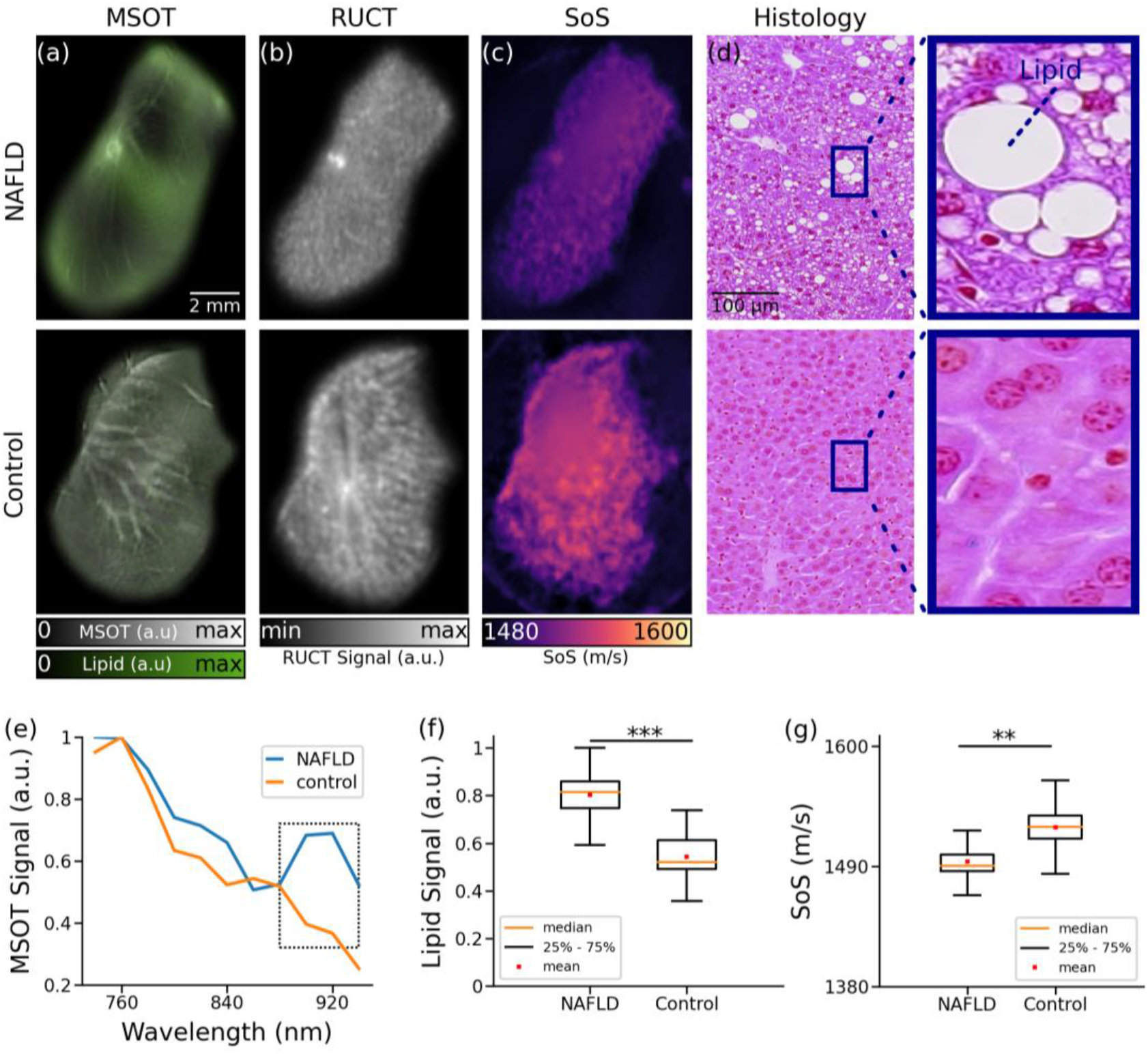
TROPUS imaging of liver tissues excised from NAFLD and control mice. (a) Un-mixed lipid distribution (green color) overlaid onto the anatomical MSOT images corresponding to averaged signals over all the acquired wavelengths for excised livers from mouse with NAFLD and control mouse. (b) Reflection ultrasound computed tomography (RUCT) images of excised livers from mouse with NAFLD and control mouse. (c) Speed of sound (SoS) images of excised livers from mouse with NAFLD and control mouse. (d) Histology images of excised livers from mouse with NAFLD and control mouse. (e) MSOT signal spectra of liver tissue shown in the panels a. (f) Average lipid signal intensities from 3 NAFLD and 3 control mice. (g) Average SoS values from 3 NAFLD and 3 control mice. (p values are indicated by * ≤ 0.05, **≤0.01 and ***≤0.001)

TROPUS was then used to image NAFLD and control mice *in vivo*. The good anatomical contrast provided by RUCT facilitated identification of the liver cross-sections (Figs. 3a-b). The SoS images further provided sufficient contrast and resolution to differentiate the liver from other surrounding tissues (Figs. 3c-d). Segmentation of the liver was done by an experienced biologist considering both the RUCT and SoS images. This served to define binary masks to quantify differences in SoS between NAFLD and control mice. SoS values were averaged for the segmented binary masks for 4 NAFLD (20 cross-sections) and 4 control (20 cross-sections) mice. Statistical analysis revealed a significant drop in SoS in liver ROIs for the NAFLD (average: 1475 m/s, STD: 34 m/s) versus control (average: 1538 m/s, STD: 18 m/s) mice (Fig. 3e, p=0.007), which is consistent with reduced SoS values in fat tissues versus healthy liver tissues [29]. A clear difference between the body weights of NAFLD and control mice was further observed (Fig. 3f, p=0.0005), with mean values of 42g and 30g, respectively. The cross-sectional areas were further calculated by manually segmenting the outer boundaries of the mouse body in the RUCT images, where the skin surface was clearly distinguishable. Statistically significant differences in cross-sectional areas of NAFLD (average area: 562 mm^2^, STD: 29 mm^2^) and control (average area: 333 mm^2^, STD: 21 mm^2^) mice were also observed (Fig. 3g, p=7e-9). Despite the increased body weight and cross-sectional area in NAFLD mice, RUCT manifested sufficient penetration depth to visualize structures in the central region of the mouse. Also, the transmitted US waves through mouse body were shown to have sufficient amplitude to enable reconstructing SoS images through the whole mouse body using the FWI reconstruction algorithm.

**Figure 3:**
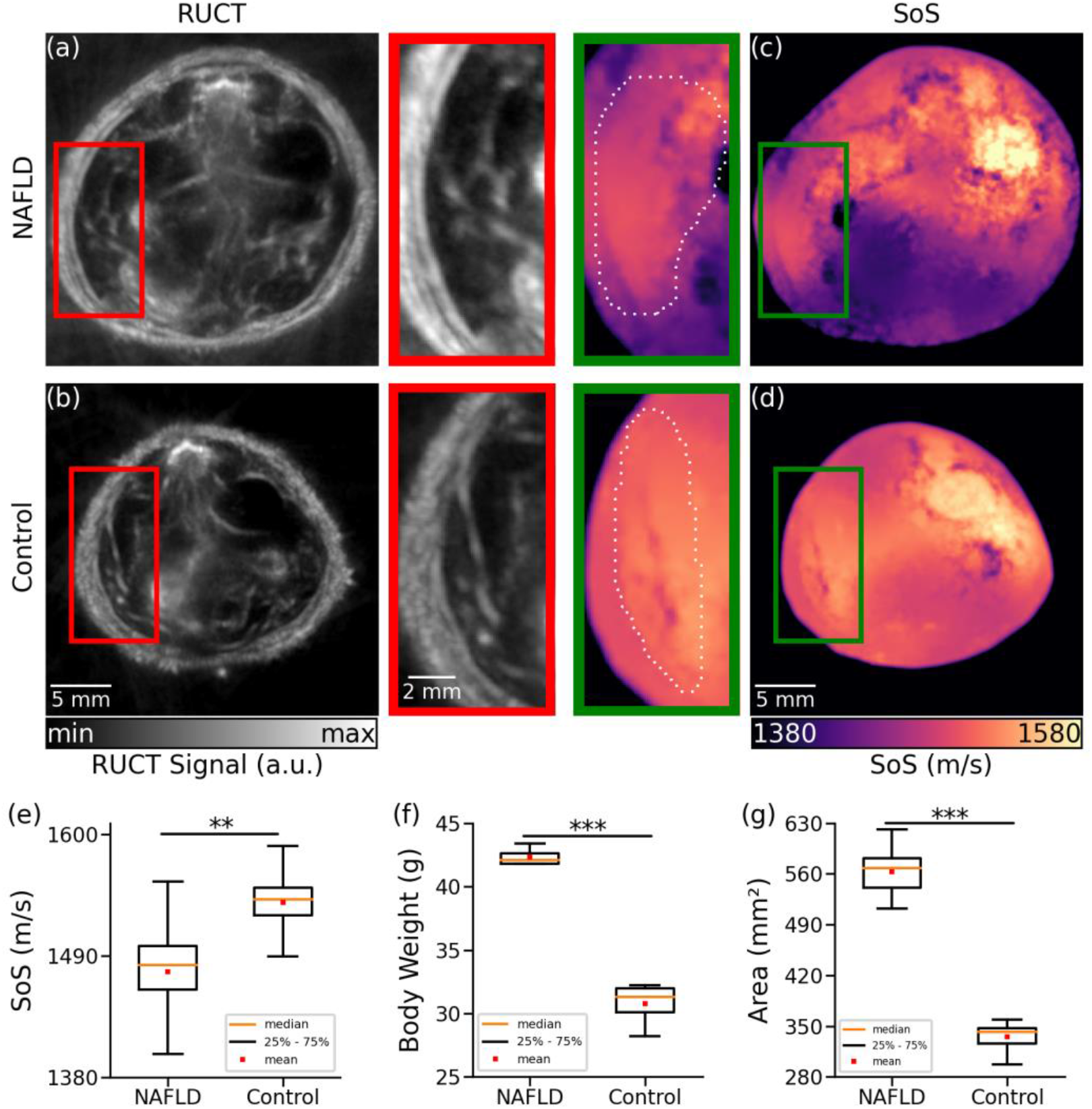
Cross-sectional reflection ultrasound computed tomography (RUCT) and speed of sound (SoS) images of NAFLD and control mice *in vivo*. (a) RUCT image of a NAFLD mouse cross section. Zoom-in of the liver region is shown. (b) RUCT image of a control mouse cross section. Zoom-in of the liver region is shown. (c) SoS image of a NAFLD mouse cross section. Zoom-in of the liver region is shown. (d) SoS image of a control mouse cross section. Zoom-in of the liver region is shown. (e) Boxplots of the measured SoS values in the segmented liver regions for NAFLD vs control mice cross sections. (f) Boxplots of the measured body weights for NAFLD and control mice. (g) Boxplots of the measured cross-sectional areas for NAFLD and control mice. (p values are indicated by * ≤ 0.05, **≤0.01 and ***≤0.001)

MSOT images were subsequently analyzed to visualize the distribution of different tissue chromophores. Specifically, linear un-mixing was performed by considering four components, namely, Hb, HbO_2_, melanin and lipids. One NAFLD and one control animal were excluded from the MSOT data analysis due to the saturated signal intensity from the melanin channel as a result of skin pigmentation. MSOT images of NAFLD and control mice corresponding to averaged signals over all the wavelengths used for acquisition are shown in Figs. 4a-b. NAFLD mice clearly manifest an increased lipid content in the liver region (Fig. 4c), indicated by the yellow contour. On the contrary, a relatively low accumulation of fat in the liver was observed in control mice (Fig. 4d). Much like for the *ex vivo* samples, analysis of the MSOT signal spectra averaged over the liver areas enabled detection of lipids. While spectra from both NAFLD and control mice monotonically decrease with wavelength, the lipid peak at 920 nm can only be detected in NAFLD mice (Fig. 4e). A statistically significant (19%) difference in lipid accumulation between NAFLD and control mice was found by calculating the averaged lipid signal values in the liver regions from all the measured cross sections (Fig. 4f, p=0.05).

**Figure 4:**
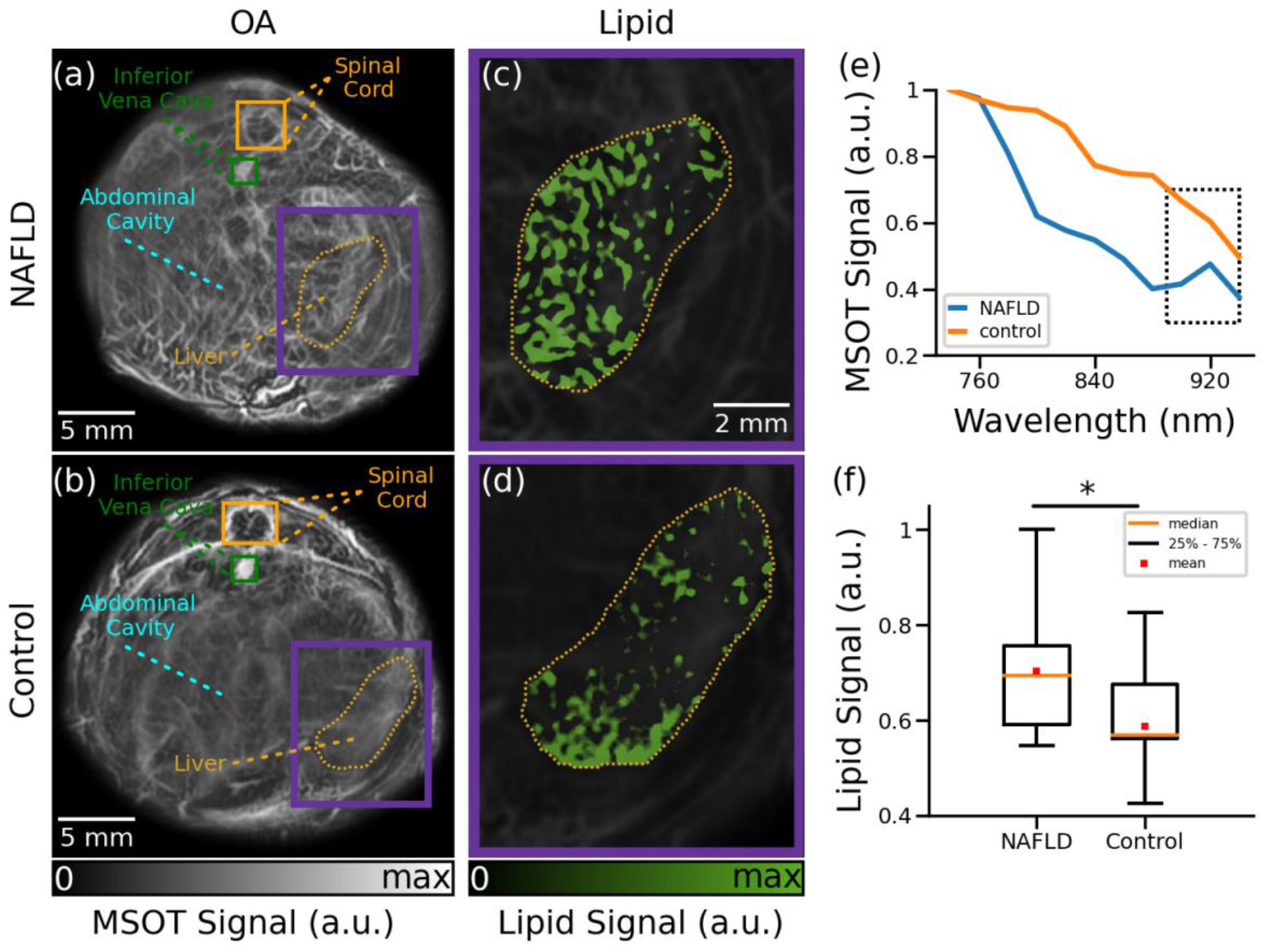
Cross-sectional MSOT images of NAFLD and control mice *in vivo*. (a) Cross section of a NAFLD mouse. (b) Cross section of a control mouse. (c) The un-mixed bio-distribution of lipids within selected liver region is shown for NAFLD mouse. (d) The un-mixed bio-distribution of lipids is shown for control mouse. (e) Spectrum of the MSOT signals in liver region indicated in panels (a-b). (f) Boxplots of the lipid signals in the liver cross sections for all mice. (p values are indicated by * ≤ 0.05, **≤0.01 and ***≤0.001)

## III. DISCUSSION

Early detection of NAFLD is essential for preventing progression of this condition to more advanced stages [31]. The process of fat accumulation in the liver is generally reversible before the onset of fibrosis by adjusting the daily diet and following a healthy lifestyle [32]. However, development of effective treatment strategies for NAFLD implies the *in vivo* validation of potential therapies in preclinical disease models. Histopathology imaging is conventionally used for this purpose [33], which however only allows measurements at single time points thus hampering longitudinal treatment follow-up studies. *In vivo* imaging modalities have thus been attempted for liver screening, predominantly MRI and pulse-echo US [34], which however suffer from low sensitivity and insufficient quantification accuracy. Multi-modal imaging with TROPUS represents a valuable alternative that can provide multi-parametric readings of the liver tissue condition. We have shown that the optical absorption peak of lipids at 920 nm facilitates quantification of fat accumulation in the MSOT images. The tomographic RUCT imaging was further shown to achieve improved resolution and contrast with respect to standard pulse-echo US, thus enabling clear delineation of the outer boundary and internal structures in the cross-sectional images. The FWI reconstruction method further enabled the rendering of accurate SoS maps. Segmentation accuracy of the liver in the images benefits from the combination between SoS and RUCT images, which facilitated the observation of statistically significant differences in the measured parameters in NAFLD versus control liver tissues.

While TROPUS exploits the synergistic combination of three modalities for detecting and evaluating liver abnormalities, each modality is associated with certain limitations. For instance, MSOT imaging is affected by light attenuation in biological tissues, which effectively limits the achievable depth [40]. The so-called spectral-coloring effects may further hamper accurate quantification of chromophore distribution in deep tissues [41]. Model-based reconstruction algorithms can be used for rendering more quantitative results, although accurate modelling of all the factors affecting MSOT signals remains challenging [37]–[40]. On the other hand, due to the need for image compounding, RUCT and SoS imaging have an inferior imaging speed as compared to MSOT. While the achieved frame rates were generally sufficient for the experiments performed in this work, a high temporal resolution may be required to quantify dynamic biological processes, such as contrast-enhanced imaging for better assessment of NAFLD. The frame rate of all the TROPUS modalities can be increased by compressed sensing methods [26], [41] or optimized data acquisition strategies [25]. Deep learning-based methods could further enhance the performance of sparse acquisition strategies and provide more accurate segmentation of the liver boundaries [42], [43]. It is also important to take into account that 3D imaging can only be achieved by vertically scanning the transducer array [24]. Alternatively, 3D imaging systems have been proposed for both MSOT and US imaging [44], [45].

Going forward, longitudinal studies starting from the early fat accumulation to fibrosis development, all the way to advanced pathological stages, such as liver cirrhosis or hepatocellular carcinoma, can also be performed with TROPUS, thus revealing new insights on the underlying mechanism of disease progression. Contrast agents may also be used for an enhanced TROPUS performance. For example, indocyanine green (ICG) is mainly cleared through the liver, and thus can be used to assess functional differences between healthy and NAFLD mice by comparing clearance time [22]. Different types of nanoparticles can also be used to boost imaging sensitivity and contrast [46]–[49]. Additional functional parameters such as blood flow can be extracted with contrast-enhanced MSOT imaging and Doppler US [50], [51].

In conclusion, we have demonstrated the capacity of the multi-modal TROPUS imaging for detecting and assessing NAFLD. The proposed approach facilitates quantification of liver abnormalities at early stages using a variety of optical and acoustic contrasts. It thus defines new *in vivo* imaging biomarkers to evaluate the efficacy of potential treatment strategies, providing a valuable alternative to conventional methods of assessing fat accumulation in the liver. All the three imaging modalities, namely, MSOT, pulse-echo, and transmission US, have already been used in clinics [18], [52], [53] laying the groundwork for translating the TROPUS approach toward diagnosis and monitoring of NAFLD in humans.

## IV. MATERIALS and METHODS

### Imaging System

The TROPUS imaging setup contains four main components, namely a circular US transducer array, a nanosecond laser source, a DAQ and a workstation PC used for the system synchronization, data transfer, storage and processing [24]. The custom-engineered ring-shaped detector array (Imasonic Sas, Voray, France) consists of 512 individual elements operated in both transmit mode for US wave generation and in receive mode for the detection of OA, pulse-echo (reflection) and transmitted US signals (Fig. 1a). The array has 40 mm radius with the individual elements having 0.37 mm x 15 mm dimensions, interelement spacing of 0.1 mm, peak central frequency of 5 MHz and transmit/receive bandwidth of 60% at −6 dB. The array’s active surface is shaped to provide cylindrical (toroidal) focusing in the imaged (2D) plane. During the experiments, the array was connected to an electronically controlled stage system with 4 degrees of freedom (x, y, z translations and azimuthal rotation) enabling accurate positioning of the imaged mouse at the center followed by volumetric scanning along the elevational (z) dimension. The mouse and the transducer array were placed in a temperature-controlled (34°C) water tank to ensure optimal physiological conditions and uninterrupted acoustic coupling. A tunable nanosecond OPO laser (SpitLight, InnoLas Laser GmBH, Krailling, Germany) was used for the OA excitation. The laser delivers ~20 mJ per pulse energy at 25 Hz repetition rate and optical wavelength between 680 and 1200 nm tunable on a per-pulse basis. The output beam was guided through an optical fiber separated into 12 output ferules with dimensions 0.21 mm x 12.65 mm to illuminate the object from different angles with uniform fluence (CeramOptec GmBH, Bonn, Germany) and optical energy density below safety limits [54]. The output ferules were equidistantly distributed on the top and bottom parts of the array with 24° tilt angle to optimize the uniformity of the illumination profile in the imaging plane. The MSOT and US signals collected by the array were digitized with a custom engineered DAQ (Falkenstein Mikrosysteme GmbH, Taufkirchen, Germany). The DAQ is connected to a workstation PC via 1 Gbit/s Ethernet to transfer the acquired signals. The workstation employed has 128 GB random access memory (RAM) and an NVIDIA GeForce GTX 1060 graphical processing unit (GPU) for real-time reconstruction of images. This PC was also used for synchronizing the delays between laser emission and US transmission, controlling the stages, and storing the acquired signals.

### Multispectral Optoacoustic Tomography (MSOT) Imaging

Imaging in the MSOT mode was performed by quickly switching the optical wavelength of the nanosecond OPO laser from 740 to 940 nm with 20 nm step size at 25 Hz repetition rate. For each laser pulse, the OA signals recorded by all 512 elements were simultaneously sampled by the DAQ at 40 megasamples per second (2030 samples were acquired per laser pulse from each element). The acquired signals were first bandpass filtered with cut-off frequencies 0.1 and 6 MHz. Then, MSOT images were reconstructed with a back-projection algorithm assigning different SoS values for the background (water) and the mouse body (Fig. 1c) [55]. 200 frames were averaged for each cross section to improve contrast-to-noise ratio (CNR). Frames affected by breathing motion were separated before averaging using an automatic detection algorithm based on the cross correlation between the frames [56]. The outer boundaries of each cross section were manually segmented in the reconstructed images using the combined information from RUCT and MSOT images. The segmented binary masks were also used to correct for light attenuation through the mouse using a simple modified Bessel function approximation [57]. Then, adaptive histogram equalization was applied on the MSOT images. After the histogram equalization, Frangi (vesselness) filter was used to detect vessels inside the mouse body [23]. As a final step, the segmented binary mask was applied for background suppression while combining reconstructed image and Frangi filtered image. The MSOT images acquired at the 11 wavelengths were used by a linear un-mixing algorithm [38] in order to separate Hb, HbO_2_, melanin and lipid components, the latter having a distinct peak in its absorption spectrum at 920 nm (Fig. 1b) [27].

### Reflection Ultrasound Computed Tomography (RUCT) Imaging

RUCT imaging was based on the STA image acquisition technique [25]. Data acquisition was performed by sequential transmission of a single-cycle bipolar square US wave (0.16 μs, 38 Vpp) with each array element. After each transmission event, all transducer elements were switched to receive the reflected and transmitted US waves. The acquisition scheme thus resulted in 512×512 time-resolved signals for the 512 transmission events. Cross-sectional images from single transmission events were reconstructed individually using delay and sum (DAS) algorithm [25]. It combines the information contained in 128 neighboring channels (90°) around the transmitting element to reconstruct a low-contrast RUCT image from each individual transmission event. Image compounding was subsequently performed by adding up the 512 low-contrast images, which resulted in a better image contrast owing to consolidation of the different views around the sample. The final (high contrast) RUCT images are presented on a logarithmic scale (Fig. 1d).

### Speed of Sound (SoS) Imaging

Data acquisition for SoS mapping was based on the same STA-based method described above. Quantitative images of the SoS distribution in m/s were reconstructed from the US waves that traversed the imaged object. Specifically, signals collected from 171 elements on the opposite side of each transmitting element were considered. A gradient-descent FWI algorithm was used to iteratively vary the estimation of the SoS values in the defined image grid to minimize the mean-squared error between the estimated waves and the actual measurements (Fig. 1e) [28]. In this work, 40 iterations were used in all cases.

FWI methods are able to improve resolution and contrast in the transmission US imaging mode when compared to the less precise bent-ray-based approach previously reported for TROPUS reconstructions [23]. Nevertheless, FWI is more computationally complex, and typically requires large computational times, even when employing a GPU. Therefore, in this work, we used two approaches to speed-up the reconstruction time. On the one hand, the initial estimation of the SoS mapping was obtained from the time-of-flight (TOF) of each emitter-receiver pair. Reference waveforms for each emitter-receiver pair were obtained from acquisitions in water, i.e. no sample placed within the FOV, and the transmitted signals were decomposed as the sum of scaled and time-shifted versions of the reference waveforms. The TOF values were obtained as the minimum of the time shifts obtained from this decomposition. This approach is more robust and less sensitive to noise than the conventional TOF picker algorithms [24], [58], [59]. On the other hand, the estimation of the transmitted waves for a specific SoS mapping within the iterative algorithm was obtained by sampling the space between each emitter and receiver with multiple paths using parallel computations on a GPU, and convolving the reference waveform with the estimated TOF of each path [60]. This avoids the need of using slow acoustic solvers. With the proposed method, the total SoS reconstruction time was within 5 minutes per slice.

### Ex Vivo Liver Imaging

*Ex vivo* imaging was performed to validate that the quantitative readings provided by TROPUS enable the differentiation between the diseased and normal liver. For this, *ex vivo* liver samples from 3 NAFLD and 3 control mice were imaged. The samples were embedded in a 20 mm cylindrical agarose phantom (1.3% w/v agarose powder). The same data acquisition protocol was executed as for the *in vivo* animal imaging experiments. Cross-sectional images from two different slices were acquired by vertically shifting the electronically controlled stage with a 1 mm step size.

Fat accumulation in liver tissues was further validated with tissue histological sections. Specifically, a Leica ASP300S tissue processor (Leica, Heerbrug, Switzerland) was used for paraffin embedding. Then, a microtome (Model: Microm HM 335 E, Thermo Scientific, Walldorf, Germany) was used to generate 3 μm thick tissue samples. The samples were stained by H&E and examined by an experienced liver histopathologist using a Nikon Eclipse 80i microscope (Nikon AG, Egg, Switzerland). 10x and 20x magnification images were acquired using a brightfield microscope.

### Animal Experiments

Animal housing and experiments were performed in accordance with the Swiss animal welfare laws approved by the Committee on Animal Experimentation for the Canton de Vaud, Switzerland (VD 3401.c). Mice of C57BL/6 background were housed at the department of Biomedical Sciences, University of Lausanne, Switzerland for 24 weeks under a 12 h dark/light cycle. The cages were ventilated and kept in a room with temperature and moisture controlled to 20-22 °C and 50-60%, respectively. After the first 8 weeks, half of the mice continued having *ad libitum* access to normal chow (Granovit, Switzerland; 3242.PX.F12) and water, while the other half was given *ad libitum* access to high fat diet (Envigo, Harlan Teklad, USA; Cat no TD.93075.PWD, Adjusted Calories Diet [55/fat]) with fructose and glucose included in the water (23.1 g/L d-fructose (Axonlab) + 18.9 g/L d-glucose (Axonlab)) for 16 weeks, to develop a diet-induced model of NAFLD [61]. Body weight was measured weekly with a digital balance.

4 mice with NAFLD and 4 control mice were imaged with TROPUS. Anaesthesia was induced with an initial dose of 4% isoflurane (Abbott, Cham, Switzerland) in an oxygen/air mixture (200:800 mL/min) and was maintained at 1.5% isoflurane supplied via a nose cone under normal air supply (oxygen/air 100:400 mL/min). The fur around the abdomen was shaved and depilated. The mice were vertically placed inside the water tank at the center of the ring array transducer with their head kept above the water surface by means of a custom-designed animal holder. Body temperature was maintained at 36.5°C by heating of the water using an electrical heater. For each mouse, 5 cross-sectional images from different sections corresponding to the liver region (40 images in total) were acquired by vertically shifting the US array with an electronically controlled stage.

## Authors’ Contributions

Conception and design: B. Lafci, X. L. Deán-Ben, D. Razansky

Development of methodology: B. Lafci, X. L. Deán-Ben, D. Razansky

Acquisition of data: B. Lafci, A. Hadjihambi, C. Konstantinou, L. Pellerin, X. L. Deán-Ben

Analysis of data: B. Lafci, N. C. Burton, J. L. Herraiz, X. L. Deán-Ben

Writing, review, and revision of the manuscript: B. Lafci, A. Hadjihambi, J. L. Herraiz, L. Pellerin, N.

C. Burton, X. L. Deán-Ben, D. Razansky

Study supervision: D. Razansky

## Acknowledgments

This work was supported by Swiss Data Science Center grant C19-04.

## Code Availability

Reconstruction code for delay and sum beamformer using synthetic transmit aperture can be found here: https://github.com/berkanlafci/pyruct

